# FinaleDB: a browser and database of cell-free DNA fragmentation patterns

**DOI:** 10.1101/2020.08.18.255885

**Authors:** Haizi Zheng, Michelle S Zhu, Yaping Liu

## Abstract

**Summary:** Circulating cell-free DNA (cfDNA) is a promising biomarker for the diagnosis and prognosis of many diseases, including cancer. The genome-wide non-random fragmentation patterns of cfDNA are associated with the nucleosomal protection, epigenetic environment, and gene expression in the cell types that contributed to cfDNA. However, current progress on the development of computational methods and understanding of molecular mechanisms behind cfDNA fragmentation patterns is significantly limited by the controlled-access of cfDNA whole-genome sequencing (WGS) dataset. Here, we present FinaleDB (FragmentatIoN AnaLysis of cEll-free DNA DataBase), a comprehensive database to host thousands of uniformly processed and curated de-identified cfDNA WGS datasets across different pathological conditions. Furthermore, FinaleDB comes with a fragmentation genome browser, from which users can seamlessly integrate thousands of other omics data in different cell types to experience a comprehensive view of both gene-regulatory landscape and cfDNA fragmentation patterns.

**Availability and implementation:** FinaleDB service: http://finaledb.research.cchmc.org/ FinaleDB source code: https://github.com/epifluidlab/finaledb_portal and https://github.com/epifluidlab/finaledb_workflow.

## 1 Introduction

Circulating cell-free DNA (cfDNA) in peripheral blood and urine have recently been shown as a promising biomarker for disease diagnosis and prognosis (Phallen et al., 2017; Cohen et al., 2018; Newman et al., 2014; Bettegowda et al., 2014; Liu et al., 2020; Shen et al., 2018). The fragment lengths of cfDNA are not uniform across the genome and are influenced by the local epigenetic environment and different physiological conditions (Snyder et al., 2016; Ivanov et al., 2015). High-resolution DNA fragment length profiling of cfDNA has revealed a predominant 167-base pair (bp) peak and 10-bp periodicity pattern in cfDNA fragments, which is highly correlated with local nucleosomal structure and histone modifications (Snyder et al., 2016; Ivanov et al., 2015). The size of cfDNA fragments differs according to their tissues-of-origin, including differences between fetal and maternal DNA, and between tumor and non-tumor derived DNA (Snyder et al., 2016). The cfDNA fragmentation patterns and their derived patterns from whole-genome sequencing (WGS), such as nucleosome positions, patterns near transcription start sites or transcription factor binding sites, ended position of cfDNA, fragmentation hotspots, co-fragmentation patterns, and large-scale fragmentation changes at mega-base level, offer extensive signals from the diseased tissues, as well as possible alterations from peripheral immune cell deaths, which can significantly increase the sensitivity for disease diagnosis(Snyder et al., 2016; Ulz et al., 2016; Jiang et al., 2018; Cristiano et al., 2019; Ulz et al., 2019; Sun et al., 2019; Liu et al., 2019; Zhou and Liu, 2020).

Due to the protection of genotype information from the patients, which is not needed for the fragmentation analysis, most cfDNA WGS datasets are deposited in the controlled-access repositories, such as the database of Genotypes and Phenotypes (dbGaP) and the European Genome-phenome Archive (EGA). The data access in these repositories requires special and lengthy application processes and sometimes even data transfer agreements that may take several months between the two organizations’ legal departments. Moreover, the cfDNA fragmentation patterns are inferred from the mapping locations of paired-end short-read sequencing, which are highly affected by the reads’ quality, length, and choices of the mapping strategy (Roadmap Epigenomics Consortium et al., 2015). These “batch effects” will significantly affect the downstream computational inference and data analysis. Currently, a centralized database with uniformly processed cfDNA datasets from a variety of physiological conditions is still not publicly available for the community.

To address these challenges, we developed FinaleDB, a comprehensive and interactive cfDNA fragmentation pattern genome browser and database that collected thousands of publicly available cfDNA WGS datasets (Figure 1A). We uniformly processed these data with quality controls through an in-house pipeline. We provided a web portal for the database, through which users can conveniently search, browse, visualize, and download cfDNA fragmentation datasets as needed.

**Figure 1.**
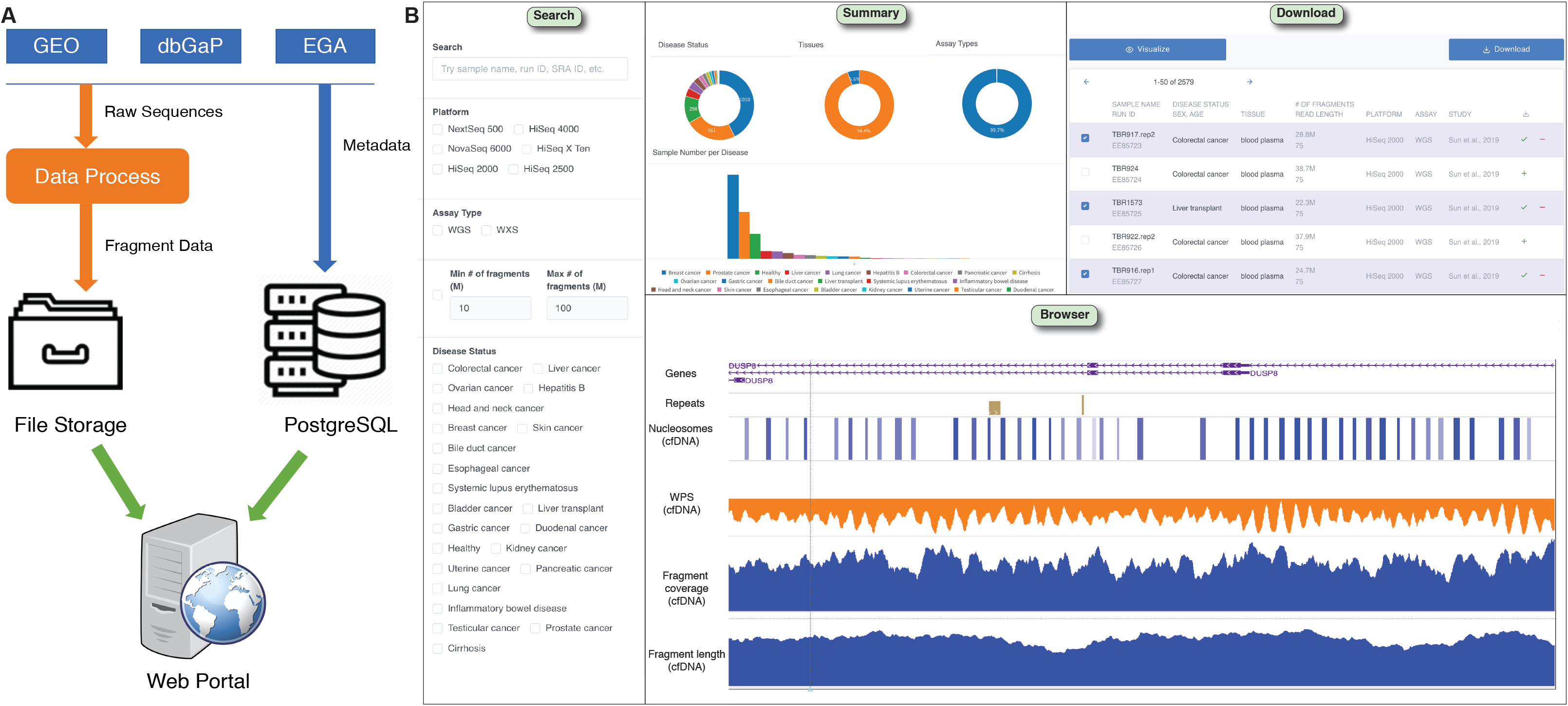
A. Overview of system design. B. Web portal and fragmentation browser.

## 2 The database

In the current version of FinaleDB, we collected 2579 paired-end cfDNA WGS datasets across 23 different pathological conditions from Gene Expression Omnibus (GEO), EGA, and dbGaP, with approximately 6TB post-processed data to download (Supplementary Table 1-2) (Snyder et al., 2016; Adalsteinsson et al., 2017; Cristiano et al., 2019; Jiang et al., 2015; Burnham et al., 2018; Rabinowitz et al., 2019; Sun et al., 2019). We processed the raw sequencing datasets by an in-house workflow (https://github.com/epifluidlab/finaledb_workflow). The workflow is managed by snakemake v5.19 and is tailored for a Kubernetes cluster with AWS Spot Instances, which ensures optimal cost-effectiveness. The cfDNA WGS datasets are collected from various studies with different read lengths. To minimize mapping bias introduced by the read lengths, we clipped all the reads from 3’end to 50bp with the additional trimming of adapters by Trimmomatic v0.39 (Bolger et al., 2014). We mapped the fragments to both hg19 and hg38 assemblies by bwa mem v0.7.17 (Li and Durbin, 2009). We removed original sequences to de-identify sensitive genotype information and only kept the high-quality fragments (both ends of reads are properly aligned with MAPQ no less than 30 and not PCR duplicated reads). For each fragment, we provided the start and end coordinates, the strand, and MAPQ score. In each dataset, we calculated the fragment coverage, the fragment length, and the Windowed-Protection Score (WPS), which reveals nucleosome positioning information (Snyder et al., 2016). Additionally, we provided the measurement of quality control (QC), such as the FastQC report, the overall distribution of fragment size, G+C% content, and library complexity.

In the back-end, we built a database powered by Amazon RDS for PostgreSQL. The database stored the essential metadata, including the sample information, sequencing platform, and study design. The fragmentation data itself is served by an HTTP static file server.

## 3 The application programming interface

The application programming interface (API) serves as an intermediate between the database and the front-end web portal. The API, based on the RESTful standard (Richardson and Ruby, 2008), can be accessed directly using any common programming language. The API currently provides seven functions (Supplementary Table 4).

## 4 The front-end web portal and fragmentation browser

We developed a web portal for the database (http://finaledb.research.cchmc.org/) based on React.js, with the source code publicly available at (https://github.com/epifluidlab/finaledb_portal) (Figure 1B). At the query page, users can search with a number of criteria, including GEO/dbGaP/EGA ID, pathological condition, source of tissue (blood or urine), associated research project, number of fragments, and sequencing platform. The visualization page comes with a modified WashU Epigenome Browser embedded within (Zhou and Wang, 2012). Users can visualize fragmentation pattern tracks of selected datasets, along with any other tracks that can be either local, remote, or those natively provided by WashU Epigenome Browser. Moreover, we offered aggregated plots of QC data for each sample chosen in the web portal, such as the overall fragment size distribution and G+C% content.

## 5 Details on methods

### 1. Data collection

In the current version, we chose 7 publications as the source of our cfDNA fragmentation data, from which we collected a total of 2579 raw paired-end sequencing datasets, as summarized below in Table 1:

**Table 1.** Publications from which the cfDNA fragment data are collected.

| Publication | Count of datasets |
| --- | --- |
| Adalsteinsson, Viktor A., et al. "Scalable whole-exome sequencing of cell-free DNA reveals high concordance with metastatic tumors." *Nature communications* 8.1 (2017): 1-13. | 1508 |
| Cristiano, Stephen, et al. "Genome-wide cell-free DNA fragmentation in patients with cancer." *Nature* 570.7761 (2019): 385-389. | 540 |
| Jiang, Peiyong, et al. "Lengthening and shortening of plasma DNA in hepatocellular carcinoma patients." *Proceedings of the National Academy of Sciences* 112.11 (2015): E1317-E1325. | 225 |
| Burnham, Philip, et al. "Urinary cell-free DNA is a versatile analyte for monitoring infections of the urinary tract." *Nature communications* 9.1 (2018): 1-10. | 141 |
| Rabinowitz, Tom, et al. "Bayesian-based noninvasive prenatal diagnosis of single-gene disorders." *Genome research* 29.3 (2019): 428-438 | 74 |
| Snyder, Matthew W., et al. "Cell-free DNA comprises an in vivo nucleosome footprint that informs its tissues-of-origin." *Cell* 164.1-2 (2016): 57-68. | 60 |
| Sun, Kun, et al. "Orientation-aware plasma cell-free DNA fragmentation analysis in open chromatin regions informs tissue of origin." *Genome research* 29.3 (2019): 418-427. | 31 |
| Total | 2579 |

We downloaded the raw sequence files (FASTQ) from the National Center for Biotechnology Information (NCBI)’s Gene Expression Omnibus (GEO) and the European Bioinformatics Institute (EMBL-EBI)’s European Genome-Phenome Archive (EGA).

The samples cover a wide range of 23 different pathological conditions (Table 2).

**Table 2.** Pathological conditions covered by FinaleDB datasets.

| Pathological condition | Count of sequencing records |
| --- | --- |
| Breast cancer | 1010 |
| Prostate cancer | 561 |
| Healthy | 298 |
| Liver cancer | 93 |
| Lung cancer | 88 |
| Hepatitis B | 67 |
| Colorectal cancer | 48 |
| Pancreatic cancer | 43 |
| Cirrhosis | 36 |
| Ovarian cancer | 30 |
| Gastric cancer | 27 |
| Bile duct cancer | 25 |
| Liver transplant | 13 |
| Systemic lupus erythematosus | 4 |
| Inflammatory bowel disease | 4 |
| Esophageal cancer | 2 |
| Testicular cancer | 2 |
| Skin cancer | 2 |
| Bladder cancer | 2 |
| Uterine cancer | 2 |
| Kidney cancer | 2 |
| Head and neck cancer | 2 |
| Duodenal cancer | 1 |
| Grand Total | 2362 |

### 2. Data process

We developed the data process pipeline using snakemake v5.19 (https://snakemake.readthedocs.io/en/stable/). The pipeline is open sourced at https://github.com/epifluidlab/finaledb_workflow.

The whole workflow is demonstrated as below in Figure 2:

**Figure 2.**
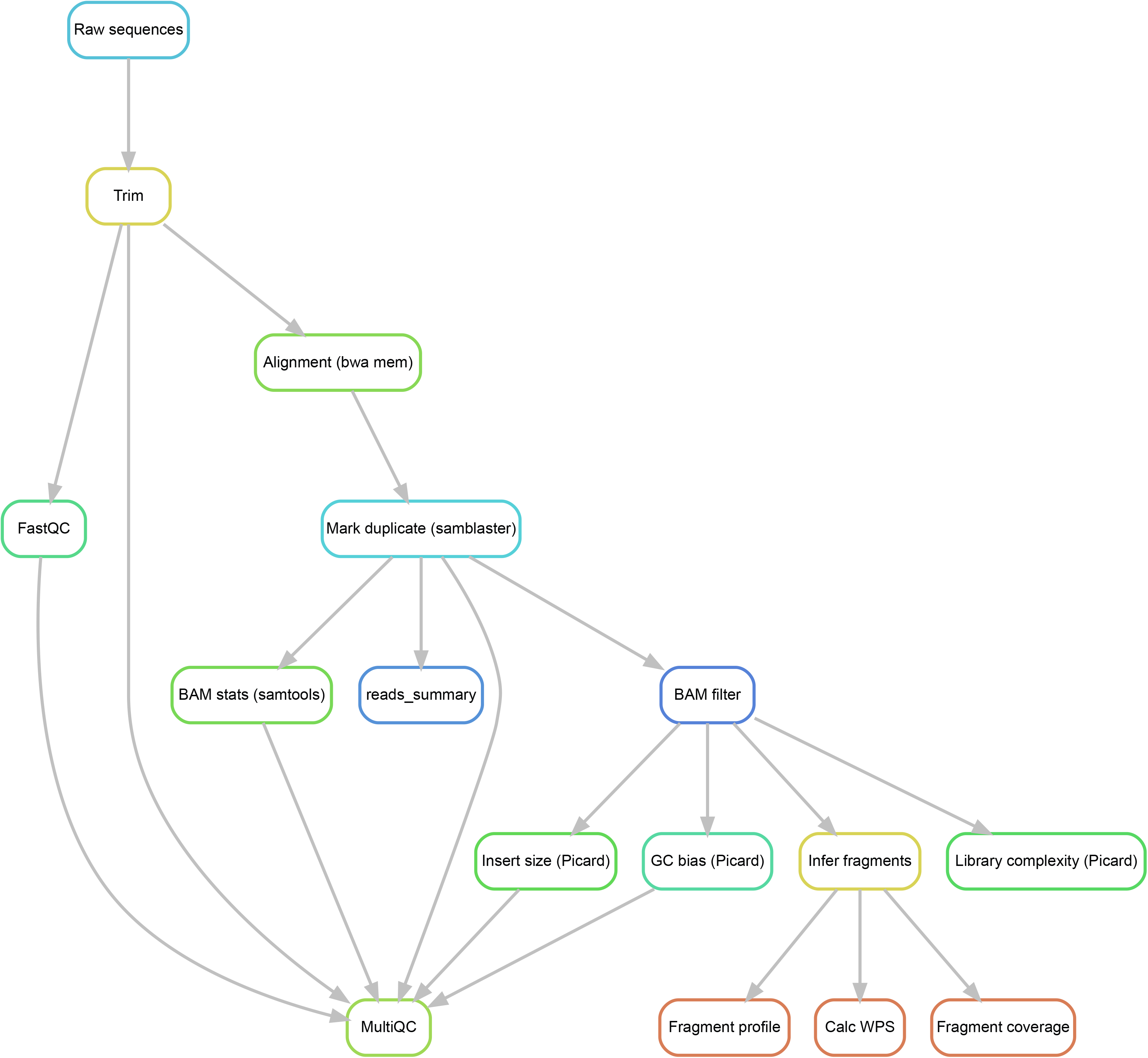
Overview of the data process workflow.

#### 2.1. Trim

We used trimmomatic v0.39 (http://www.usadellab.org/cms/?page=trimmomatic) to perform quality trim and Illumina adaptor clipping. It also crops all reads to 50bp, in order to reduce potential batch effects.

The arguments we used:

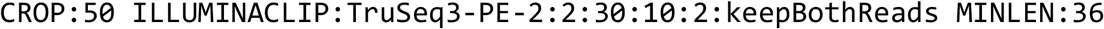

The trimmomatic adapter file we used: https://github.com/timflutre/trimmomatic/blob/master/adapters/TruSeq3-PE-2.fa

#### 2.2. Alignment

We used bwa mem v0.7.17 (https://github.com/lh3/bwa) to perform the reads alignment, with the following reference genome assemblies:

- For hg19: ftp://ftp-trace.ncbi.nih.gov/1000genomes/ftp/technical/reference/human_g1k_v37.fasta.gz
- For hg38: ftp://ftp.ncbi.nlm.nih.gov/genomes/all/GCA/000/001/405/GCA_000001405.15_GRCh38/seqs_for_alignment_pipelines.ucsc_ids/GCA_000001405.15_GRCh38_no_alt_analysis_set.fna.gz

#### 2.3. Mark duplicates

We used samblaster v0.1.25 (https://github.com/GregoryFaust/samblaster/) to mark PCR duplicates. Compared with other duplicate marking tools such as Picard MarkDuplicates, samblaster performs in a single-pass manner and thus is more effective in certain scenarios.

Entries in the BAM file are marked as duplicates will further be removed in the BAM fiter step.

#### 2.4. BAM filter

In order to perform downstream analysis including GC bias, fragment inference, we first filtered the BAM file with the following argument

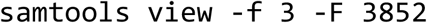

This filter rule excluded any reads which:

- Are not properly paired, or
- Are secondary alignments, or
- Failed platform/vendor quality checks, or
- Are PCR or optical duplicate (marked duplicates), or
- Are part of a chimeric alignment (supplementary alignment)

#### 2.5. GC bias

We used Picard Tools v2.22.3 (https://github.com/broadinstitute/picard/) to perform the GC bias analysis, with the following arguments:

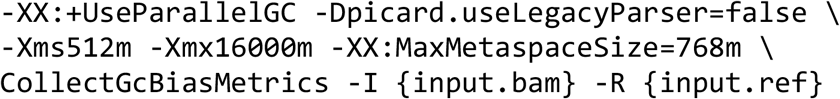

Here, `{input.ref}` indicates the reference genome used in the corresponding alignment.

#### 2.6. Library complexity estimation

We used Picard Tools v2.22.3 (https://github.com/broadinstitute/picard/) to estimate the library complexity, which is defined as the number of unique fragments in the library, with PCR duplicates excluded. A library with low complexity may indicate high PCR duplicate ratio and may negatively affect the downstream analysis.

We estimated the library complexity with the following arguments:

We used Picard Tools v2.22.3 (https://github.com/broadinstitute/picard/) to perform the GC bias analysis, with the following argument:

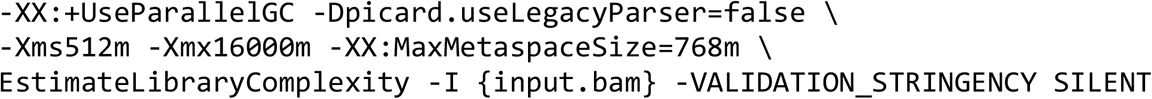

#### 2.7. Fragment inference

From each filtered BAM file, we extracted the outer fragment endpoint coordinates for each pair of aligned reads. The insert between the paired endpoints are recognized as a fragment, and the interval is reckoned as the fragment length.

The fragment data is stored in a tab-separated value (TSV) file, with <u>the same column definition as</u> BedGraph format. Each fragment data file contains five columns (Table 3):

**Table 3.**
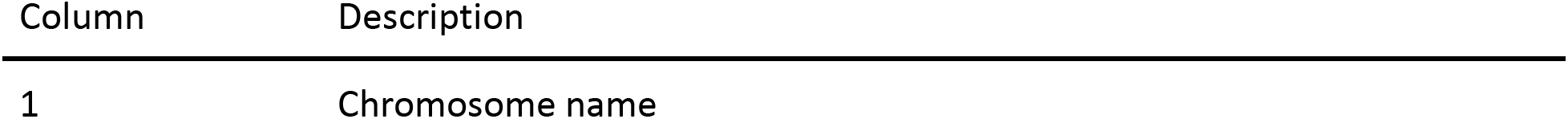
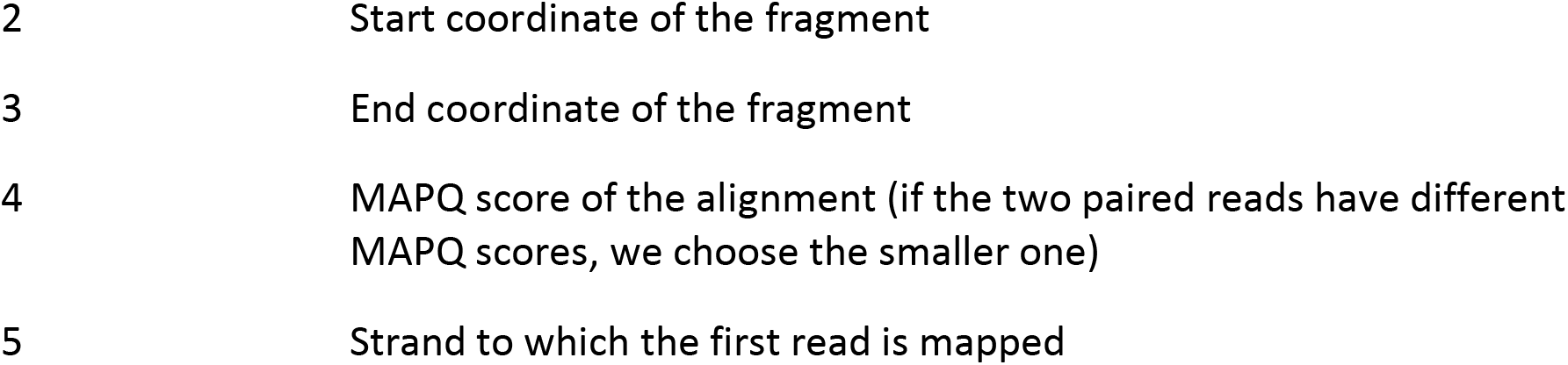
The columns in the fragment data file.

| Column | Description |
| --- | --- |
| 1 | Chromosome name |
| 2 | Start coordinate of the fragment |
| 3 | End coordinate of the fragment |
| 4 | MAPQ score of the alignment (if the two paired reads have different MAPQ scores, we choose the smaller one) |
| 5 | Strand to which the first read is mapped |

We choose the UCSC 0-based half-open convention for the coordinate system.

#### 2.8. Fragment coverage profile

To calculate the fragment coverage, we first ruled out fragments with MAPQ scores less than 30, which either indicate low mapping qualities, or reads not uniquely mapped to the reference genome (when MAPQ equals 0).

We then defined a fragment’s coverage as all positions that lie between its two ends, with the endpoints included.

#### 2.9. Fragment length profile

Similar to the fragment coverage calculation, we excluded all fragments with MAP scores less than 30.

The fragment length is defined as the interval between fragment ends, with both endpoints included, that is, the difference between Column 2 and Column 3 in the fragment data file.

#### 2.10. Windowed Protection Score (WPS)

Similar to the fragment coverage calculation, we excluded all fragments with MAP scores less than 30. Moreover, we exclude all fragments that are shorter than 120bp or longer than 180bp.

For a given coordinate in the genome, we first defined a 120bp window symmetric to the said coordinate. We then calculated the WPS which is defined as the number of fragments completely spanning the window, minus the number of fragments with one of their endpoints located within the same window.

### 3. Web portal manual

#### 3.1. Query

To the left of the page is the filter pane, where users can make database queries. In the “Search” text box, users can search by sample names (e.g. MBC_1292), sequencing run ID (e.g. EE86284), or SRA ID (e.g. SRR6708248).

In addition to the search box, users can also search through the facet filter described below,

- Platform: the sequencer used to generate the data
- Assay type: currently there are two assay types in the database: whole genome sequencing (WGS) and whole exome sequencing (WES)
- Min/max # of fragments: specify a range and only return the records with the number of fragments within
- Disease status: return records associated with any particular pathological conditions
- Tissue: return records associated with a particular tissue type. Currently, there are two tissue types: blood plasma and urine
- Publication: return records related to a particular published study

To the right of the page is the sample table, where users can browse the query results and perform the following actions:

- Click the checkbox to the left of each record to select/unselect, and then click the “Visualize” button to further inspect the selected records.
- Click the green plus sign to the right of each record to add them to the download list. Users can then click the “Download” button for the download page to download selected data files.

#### 3.2. Visualization

Regarding the selected records, this page will display the fragment size distribution, as well as showing the coverage profile, fragment size profile, and the WPS track.

To the upper left is the reference genome. Users can switch between hg19 and hg38 and inspect corresponding fragment data.

Another useful feature is importing sessions from external sources. WashU Epigenome Browser (http://epigenomegateway.wustl.edu/browser/) has a feature enabling its users to save a session and share the contents with other users or to save it for future reference. Leveraging such features, FinaleDB can integrate active tracks from the external browser to the visualization page, allowing users to view cfDNA fragmentation pattern tracks and other external tracks side by side.

To do so, first go to WashU Epigenome Browser and load any tracks of interest, then click “Session” under the “Apps” tab, and download the session file. Next, return to the FinaleDB visualization page, and upload the session file either by clicking the “Import session” zone or drag-n-drop.

#### 3.3. Download

In the download center, users can make finer selections about which files to download, and launch the download process. Currently, FinaleDB provides the download access for the following data:

- Fragment data file (.tsv)
- Fragment coverage profile (bigWig)
- Fragment size profile (bigWig)
- Windowed Protection Score (<u>bigWig</u>)

Due to the nature of the data files, FinaleDB can’t on-the-fly bundle the selected files into a single compressed archive for download. Therefore, when users click the download button, all selected files will be downloaded separately. Some browsers may apply restrictions for this multi-file download behavior. In this case, please refer to the browser documentation for more information.

### 4. System design

The entire FinaleDB system consists of three parts:

1. Metadata database
2. Static file server
3. Web portal server

#### 4.1. Metadata database

This database stores all metadata information. It is a PostgreSQL database and comes with the following tables:

1. seqrun: each record represents a “sequencing run” for a specific sample. Each seqrun record is identified by a unique ID starting with “EE”, such as EE86284.
2. sample: each record contains information of a particular sample
3. publication: each record represents a publication of a specific study project

The database is implemented with Amazon Relational Database Service (RDS), which is a service provided by Amazon Web Services (AWS).

#### 4.2. Static file server

While the metadata information is stored in the PostgreSQL database described above, all fragmentation pattern analysis results themselves are stored as static files, in standard bioinformatics data formats.

To be more specific, these files include:

1. Alignments are stored as BAM files
2. Fragment files are stored as gzipped and indexed tab-separated value (TSV) files
3. Coverage, fragment profiles and WPS tracks are stored as bigWig files for genome browser visualization
4. QC reports (FastQC, Picard GC bias and library complexity estimation) are stored in plain text files

All the files are stored in an AWS S3 bucket which is publicly available over HTTPS.

#### 4.3. Web portal

The web portal is a single-page app (SPA) developed on React.js (v16, https://reactjs.org/), a popular JavaScript web development framework, as well as client-side interface libraries listed below:

- Tabler-react (http://tabler-react.com/) for UI components
- Bootstrap React (https://react-bootstrap.github.io/) for UI components
- Highcharts (https://www.highcharts.com/) for interactive charts in the visualization page
- WashU Epigenome Browser for the displaying epigenomic tracks in the visualization page

The source code for the web portal is publicly available at https://github.com/epifluidlab/finaledb_portal under MIT License.

### 5. Programmatic access

In addition to the web interface, FinaleDB provides a set of APIs for programmatic access. We believe it facilitates researchers to make the most of the database.

Below is a brief description of the API set.

#### 5.1. Database summary

Endpoint: http://finaledb.research.cchmc.org/api/v1/summary

Usage: return summarized statistics about FinaleDB database, such as the number of samples, a list of diseases present in the database, tissue types, etc.

#### 5.2. Database queries

Endpoint: http://finaledb.research.cchmc.org/api/v1/seqrun?limit=5

Usage: return sequencing runs matching the query terms specified in the query string. For example, the API request below returns all records related to (Table 4):

**Table 4.**
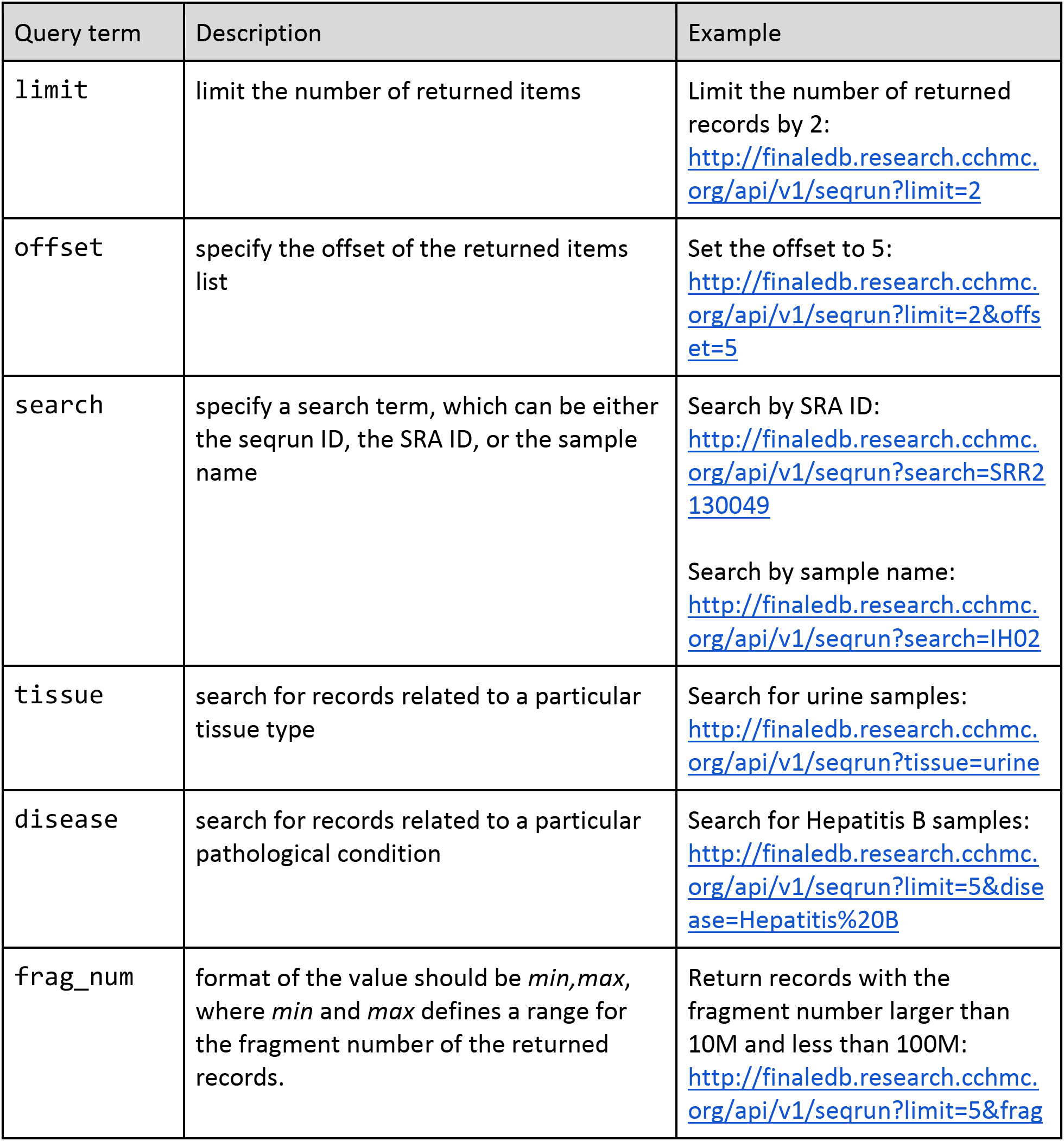
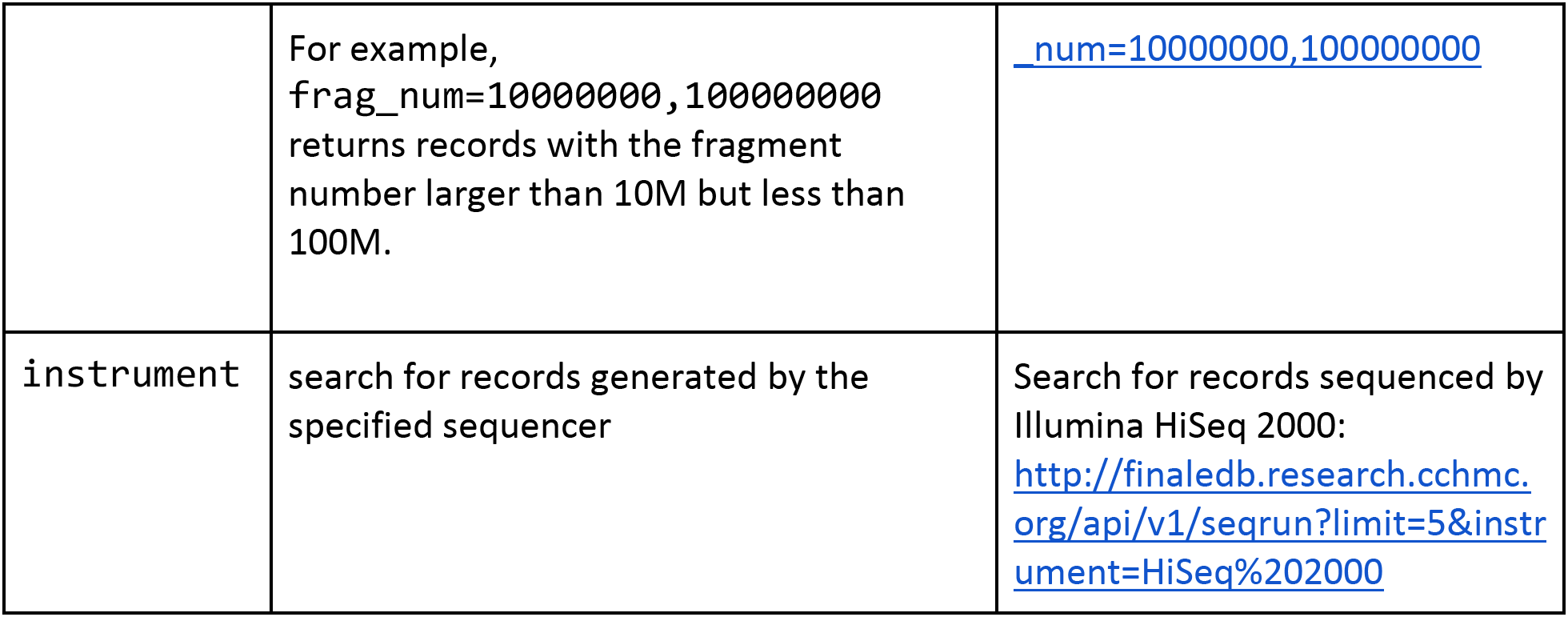
Function for RESTful API.

## Acknowledgements

The authors greatly acknowledge the supports from Biomedical Informatics (BMI) high performance computing cluster in CCHMC and Pittsburgh Supercomputing Center (PSC).

